# Helminth infection dynamics in rehabilitating Javan slow lorises are driven by time since deworming rather than host traits

**DOI:** 10.64898/2026.04.23.720522

**Authors:** Abdullah Langgeng, Marie Sigaud, Wendi Prameswari, Nur Purba Priambada, Puji Rianti, Richard Moore, Andrew J. J. MacIntosh, Ikki Matsuda

**Affiliations:** Wildlife Research Center of Kyoto University, 2□24, Tanaka□Sekiden□Cho, Sakyo, Kyoto 606□8203, Japan; Centre d’Écologie et des Sciences de la Conservation (CESCO), Muséum national d’Histoire naturelle, Centre National de la Recherche Scientifique, Sorbonne Université, Paris, France; Yayasan Inisiasi Alam Rehabilitasi Indonesia, Bogor, West Java, Indonesia; Department of Biology, Faculty of Mathematics and Science, IPB University, Bogor, West Java, Indonesia; Program of Bio-conservation, Primate Research Center, IPB University, Bogor, West Java, Indonesia; Wilder Institute, Calgary, Canada; Chubu Institute for Advanced Studies, Chubu University, 1200, Matsumoto-cho, Kasugai-shi, Aichi 487-8501, Japan; Institute for Tropical Biology and Conservation, Universiti Malaysia Sabah, Kota Kinabalu, Malaysia

**Keywords:** wildlife rehabilitation, gastrointestinal parasites, anthelmintic treatment, reinfection dynamics, parasite management, release readiness

## Abstract

Wildlife rehabilitation plays a central role in the conservation of threatened primates, yet parasite dynamics during captivity are rarely reported, particularly in relation to release readiness. We investigated gastrointestinal helminth infection patterns in rehabilitating Javan slow lorises (*Nycticebus javanicus*), a Critically Endangered species heavily impacted by the illegal wildlife trade. Using repeated fecal sampling (147 samples from 19 adults) and Bayesian mixed-effects models, we examined parasite richness, Shannon diversity, infection probability, and egg-shedding intensity in relation to release readiness status, sex, housing condition, and time since anthelmintic treatment. Four nematode taxa identifiable through egg morphology were detected: *Strongyloides* spp., strongylids, oxyurids, and *Trichuris* spp.. Parasite richness and Shannon diversity showed no credible associations with release readiness or other host and management variables. In contrast, infection probability for *Strongyloides* spp. and strongylids increased with time since deworming, and *Strongyloides* egg counts exhibited a similar temporal pattern, consistent with post-treatment reinfection dynamics. Release readiness did not predict detection probability or parasite intensity for any parasite group, despite marked differences in captivity duration and health history between individuals deemed ready for release or not. These findings indicate that gastrointestinal helminth dynamics in rehabilitating slow lorises are driven primarily by treatment-related temporal processes and individual-level heterogeneity rather than coarse host classification. They also highlight the need for longitudinal parasite monitoring and for future work evaluating how infection dynamics, management interventions, and host health relate to rehabilitation and translocation outcomes..

## INTRODUCTION

Parasites constitute dynamic components of primate ecosystems, shaping host health, survival, and population processes across both wild and managed environments. Rather than representing static indicators of host condition, parasite communities reflect ongoing ecological interactions among hosts, conspecifics, and their environments, operating across multiple temporal scales (Sanchez et al., 2018; Milotic et al., 2020). In captive and semi-captive settings such as zoological institutions and wildlife rehabilitation centers, gastrointestinal parasites are therefore of particular concern for animal welfare, disease management, and conservation planning (Schapiro and Bernacky, 2012; Li et al., 2015; Hildebrand and Zalesny, 2025). Captivity alters parasite transmission pathways by restructuring host density, contact networks, and environmental exposure, while simultaneously introducing veterinary interventions that selectively suppress some taxa but allow others to persist. As a result, parasite assemblages in managed populations often undergo qualitative reorganization rather than uniform declines in infection or diversity (Stringer and Linklater, 2014; Nunn et al., 2015; Herrera et al., 2019). Although routine anthelmintic treatment, enclosure sanitation, and controlled diets can substantially reduce infection intensity, particularly for parasites with complex life cycles requiring intermediate hosts, infections often persist for helminths with direct transmission pathways and environmentally resistant stages, which facilitate rapid reinfection under conditions of prolonged housing or repeated environmental exposure (Grove et al., 1988; Schapiro and Bernacky, 2012; Li et al., 2015). Understanding how host attributes and management practices interact to shape these dynamic parasite processes is therefore essential for effective health management and conservation-oriented decision making in rehabilitated primates.

Against this background, parasite infection dynamics can reflect ecological processes that operate over time and respond to changes in host exposure, susceptibility, and management conditions, rather than representing fixed properties of individual hosts (Altizer et al., 2006; Pedersen and Fenton, 2007). In rehabilitation contexts, parasite prevalence and intensity may vary through cycles of treatment, reinfection, and persistence, instead of converging toward stable endpoints (Grove et al., 1988; Schapiro and Bernacky, 2012; Li et al., 2015). Such temporal variability highlights that parasite infections often capture ongoing interactions between hosts and their environments, shaped by both biological and management-related processes. This sensitivity to environmental and management conditions suggests that parasites can provide valuable insights into how captive or rehabilitative settings influence biological processes over time (Sanchez et al., 2018; Herrera et al., 2019). At the same time, it cautions against interpreting parasite burdens as simple or universal indicators of host condition or release readiness (Sanchez et al., 2018), as observed infection levels may be influenced by recent interventions or exposure histories rather than intrinsic host quality alone. Addressing these dynamics requires analytical approaches that account for repeated measurements, individual heterogeneity, and time since intervention (Herrera et al., 2019), considerations that remain inconsistently incorporated in studies of parasites in rehabilitating wildlife.

Slow lorises (*Xanthoycticebus* and *Nycticebus* spp.; Nekaris and Nijman, 2022) are nocturnal strepsirrhine primates distributed across Southeast Asia and are among the most threatened primates globally due to habitat loss, fragmentation, and intensive exploitation for the illegal wildlife trade (Nekaris and Starr, 2015). The Javan slow loris (*N. javanicus*) is classified as Critically Endangered, with substantial numbers of individuals entering rescue and rehabilitation facilities following confiscation from trade (Moore et al., 2014). As a result, rehabilitation programs therefore play a central role in the conservation management of this species. Rehabilitated slow lorises often experience prolonged periods of captivity and exhibit a suite of conditions, including dental trauma, altered diets, and exposure to anthropogenic environments, that may influence parasite acquisition, persistence, and transmission dynamics (Moore et al., 2014; Rode-Margono et al., 2015). Many of these same factors also contribute to assessments of release readiness in rehabilitation programs, as dental condition, health status, and behavioral competence are key criteria used to determine whether individuals are suitable for translocation (Moore et al., 2014; Goldenberg et al., 2022). Despite this, published empirical data on parasite infections in rehabilitating slow lorises remain scarce. Most available information on helminths in *Nycticebus javanicus* comes from studies of wild populations, which have documented gastrointestinal parasites such as *Necator, Lemuricola*, and *Trichostrongylus*, but reported no association between host sex and parasite prevalence or intensity (Rode-Margono et al., 2015), leaving open questions regarding the drivers of infection heterogeneity. In parallel, molecular analyses of *Strongyloides* infecting Bornean slow lorises (*N. menagensis*) have revealed genetically distinct lineages clustering separately from *Strongyloides* species infecting other primates, suggesting cryptic diversity and potential host specificity within the genus *Nycticebus* (Frias et al., 2018). However, how parasite prevalence, abundance, and community structure vary among rehabilitating slow lorises, and whether such patterns are influenced by host traits or rehabilitation-related factors, remains poorly understood.

Building on these considerations, the present study aimed to characterize gastrointestinal helminth infections in rehabilitating *N. javanicus* housed at a wildlife rehabilitation center in Java, Indonesia. Using repeated fecal sampling and Bayesian mixed-effects models, we examined patterns of infection presence, abundance, and infracommunity diversity in relation to host traits (sex), housing condition (social versus solitary), release candidacy status, and time since anthelmintic treatment.

Because individuals differ in captivity duration, health history, and exposure to management interventions, these factors may influence parasite infection patterns during rehabilitation. We therefore evaluated whether parasite detection probability, parasite intensity, and infracommunity diversity varied with host traits, housing conditions, release candidacy status, and time since anthelmintic treatment. Given the prolonged duration of captivity experienced by many individuals, we tested whether the infection patterns differed between individuals with different rehabilitation histories. We further hypothesized that socially housed individuals would exhibit higher infection risk than solitary individuals due to increased contact rates, and that parasite prevalence and intensity would increase with time since deworming, reflecting post-treatment reinfection dynamics. By integrating host- and management-related predictors, this study sought to clarify how rehabilitation practices shape parasite dynamics in slow lorises and to inform evidence-based health management and release decision-making in conservation and reintroduction programs.

## METHODS

### Study Subjects and Sample Collection

Sample collection was conducted from June to October 2024 at the YIARI rehabilitation center in West Java, Indonesia. We studied 19 adult rehabilitating *N. javanicus*, (12 females and 7 males). Among them, ten individuals (7 females and 3 males) were categorized as release candidates (hereafter: candidates) based on veterinary and behavioral assessments conducted by the rehabilitation team, including, oral and general health, medical history, behavioral activity levels. These individuals were deemed suitable for release and were subsequently translocated to Mount Halimun-Salak National Park, West Java, by the end of October 2024. The remaining nine individuals (5 females and 4 males) were considered non-releasable (hereafter: non-candidates) and retained permanently at the center because combinations of dental impairments, recurrent or chronic health problems, and/or physical handicaps were judged to limit their long-term viability in the wild. Exploratory logistic regression models treating candidacy as the response variable indicated that rehabilitation duration, dental condition, and medical history were the strongest contributors to an individual’s candidacy status (McFadden pseudo-R^2^: 0.31 – 0.84). Release candidates had spent 5–32 months at the rehabilitation center (mean ± SD: 11.8 ± 8.9) and an additional two weeks in a soft-release enclosure before being released into Mount Halimun-Salak National Park, West Java. Non-candidates had resided at the center for substantially longer periods, ranging from 29 to 145 months (mean ± SD: 84.4 ± 35.0). All individuals received anthelmintic treatment with albendazole (10 mg/kg body weight). Treatments were administered to all individuals on 28 December 2023, roughly 6 months prior to the start of this study, followed by a subsequent treatment on 1 September 2024 for non-candidates and 15 October 2024 for candidates prior to soft release.

Individuals were sampled repeatedly throughout the study period, yielding 147 fecal samples from 19 individuals (mean ± SD = 7.7 ± 3.4 samples per individual, range = 4–14). Samples were collected opportunistically within one hour of defecation and preserved in sodium acetate-acetic acid-formaldehyde (SAF) solution at a 3:1 SAF-to-sample ratio for coproscopic analysis. Fecal samples were processed using a modified formalin-ethyl acetate sedimentation protocol to concentrate helminth eggs (MacIntosh et al., 2010). Egg counts were determined microscopically using a McMaster chamber, and results were expressed as eggs per gram of fecal sediment (EPG), which served as a proxy for parasite infection intensity across individuals. Parasite eggs were identified based on morphological characteristics and size (Modrý et al., 2018). Although EPG does not represent true adult worm burdens, it is a standard and widely used index of parasite infection in wildlife parasitology, particularly when direct quantification of parasites is not feasible (Gillespie, 2006; Modrý et al., 2018). Because the relationship between EPG and actual worm counts is unknown in the present study system, results should be interpreted with appropriate caution.

### Data Analysis

We used three complementary metrics to characterize gastrointestinal parasite infections in focal subjects following standard definitions in parasite ecology (Bush et al., 1997). First, parasite alpha diversity was quantified at the host individual level based on parasite species richness and Shannon diversity. Second, detection probability for each parasite taxon was modeled using presence–absence of parasite eggs per fecal sample as the response variable. Third, parasite infection intensity was quantified as eggs per gram of feces (EPG) and used as a proxy for infection intensity.

To test predictions regarding variation in parasite detection probability and intensity, we fitted Bayesian generalized linear mixed-effects models using the *brms* package in R (Bürkner, 2017). To evaluate the effects of rehabilitation-related factors on gastrointestinal parasite infections, we included release candidacy status, sex, housing type (social versus solitary), and weeks post-deworming (mean ± SD: 26.96 ± 12.59; range: 1–39; median: 32) as fixed effects. To account for repeated sampling and avoid pseudoreplication, individual identity and sampling date were included as random intercepts.

Parasite egg counts (EPG; rounded to the nearest integer) were initially modeled using negative binomial distributions to account for overdispersion in count data. Because egg counts also contained many zero values, we additionally evaluated zero-inflated negative binomial (ZINB) models to account for potential excess zeros. Standard negative binomial and ZINB models were compared using leave-one-out cross-validation (ELPD-LOO), and the model with stronger predictive support was retained for inference. Parasite presence or absence was modeled as a binary response using Bernoulli distributions with a logit link. Weakly informative priors were applied to all model parameters to stabilize estimation under sparse data conditions. For Bernoulli models of infection presence– absence, fixed effects were assigned Normal (0, 2.5) priors on the logit scale, the intercept was assigned a Student-t (3, 0, 2.5) prior, and random-effect standard deviations were assigned Exponential(1) priors. For parasite egg count models, fixed effects were assigned Normal (0, 1) priors on the log scale and the negative binomial shape parameter an Exponential (1) prior. In ZINB models, the zero-inflation probability was assigned a Normal (−1.5, 1) prior on the logit scale. Prior predictive checks were conducted to ensure that the specified priors generated biologically plausible parasite count and prevalence distributions. Pareto-k diagnostics were used to assess the reliability of LOO estimates. Additional diagnostic results are provided in the Supplementary Materials (Tables S2, S3, andS5; Figures S1–S3)

## RESULTS

We detected at least four helminth taxa (Figure S4) in 147 fecal samples collected from 19 individual *N. javanicus*. At the individual level, *Strongyloides* spp. was the most prevalent parasite, detected in 84.2% of sampled individuals, followed by strongylids (36.8%), oxyurids (31.6%), and *Trichuris* spp. (15.8%). At the sample level, mean egg counts were highest for *Strongyloides* spp. (169.80 ± 362.22 EPG; range: 0–2381), followed by oxyurids (26.67 ± 84.40 EPG; range: 0–601), strongylids (20.82 ± 48.82 EPG; range: 0– 226), and *Trichuris* spp. (20.22 ± 103.93 EPG; range: 0–1139).

### Richness and Shannon Diversity

At the sample level, parasite richness averaged 0.98 ± 0.94 taxa (range: 0–4), and Shannon diversity averaged 0.17 ± 0.29 (range: 0–1.10), indicating generally low parasite diversity per fecal sample. Neither parasite richness nor Shannon diversity showed strong associations with release candidacy or other rehabilitation-related variables, although both metrics tended to increase with time since deworming (Table 1).

**Table 1.** Bayesian mixed-effects model results for gastrointestinal parasite alpha diversity (Shannon diversity and richness) in rehabilitating *N. javanicus*. Posterior means (β) and 95% credible intervals (CrI).

| Predictor | Shannon diversity | Parasite richness |
| --- | --- | --- |
| **Fixed effects** |  |  |
| Intercept | 0.19 (−0.05, 0.45) | −0.27 (−1.08, 0.46) |
| Status (non-candidate) | 0.10 (−0.33, 0.52) | 0.45 (−0.76, 1.56) |
| Sex (male) | −0.04 (−0.29, 0.20) | 0.16 (−0.59, 0.91) |
| Housing (solitary) | −0.06 (−0.37, 0.25) | −0.33 (−1.23, 0.61) |
| Rehabilitation duration | 0.11 (−0.07, 0.30) | 0.35 (−0.15, 0.90) |
| Week post-deworming | 0.02 (−0.01, 0.06) | 0.18 (−0.01, 0.38) |
| **Random effects (SD)** |  |  |
| Host ID | 0.24 (0.15–0.36) | 0.66 (0.35–1.14) |

### Detection probability

Bayesian mixed-effects logistic regression models indicated that, for a subset of parasite taxa, detection probability was associated with time since anthelmintic treatment. Specifically, detection probability increased with weeks post-treatment for strongylids and *Strongyloides* spp. In addition, *Strongyloides* egg counts increased with weeks post-treatment in the abundance models. These credible temporal effects are summarized in Figure 1. In contrast, no strong evidence was detected for effects of release status (candidate vs. non-candidate), sex, or housing condition on infection probability for any parasite group, including *Trichuris* spp. and oxyurids, as all corresponding credible intervals overlapped zero (Table 2). Rehabilitation duration also showed consistently positive posterior associations with detection probability across several parasite taxa. Although 95% credible intervals overlapped zero, effect sizes on the odds scale were substantial, corresponding to approximately 3.6–4.9-fold increases in detection odds per 1 SD increase in rehabilitation duration (Table S4).

**Table 2.**
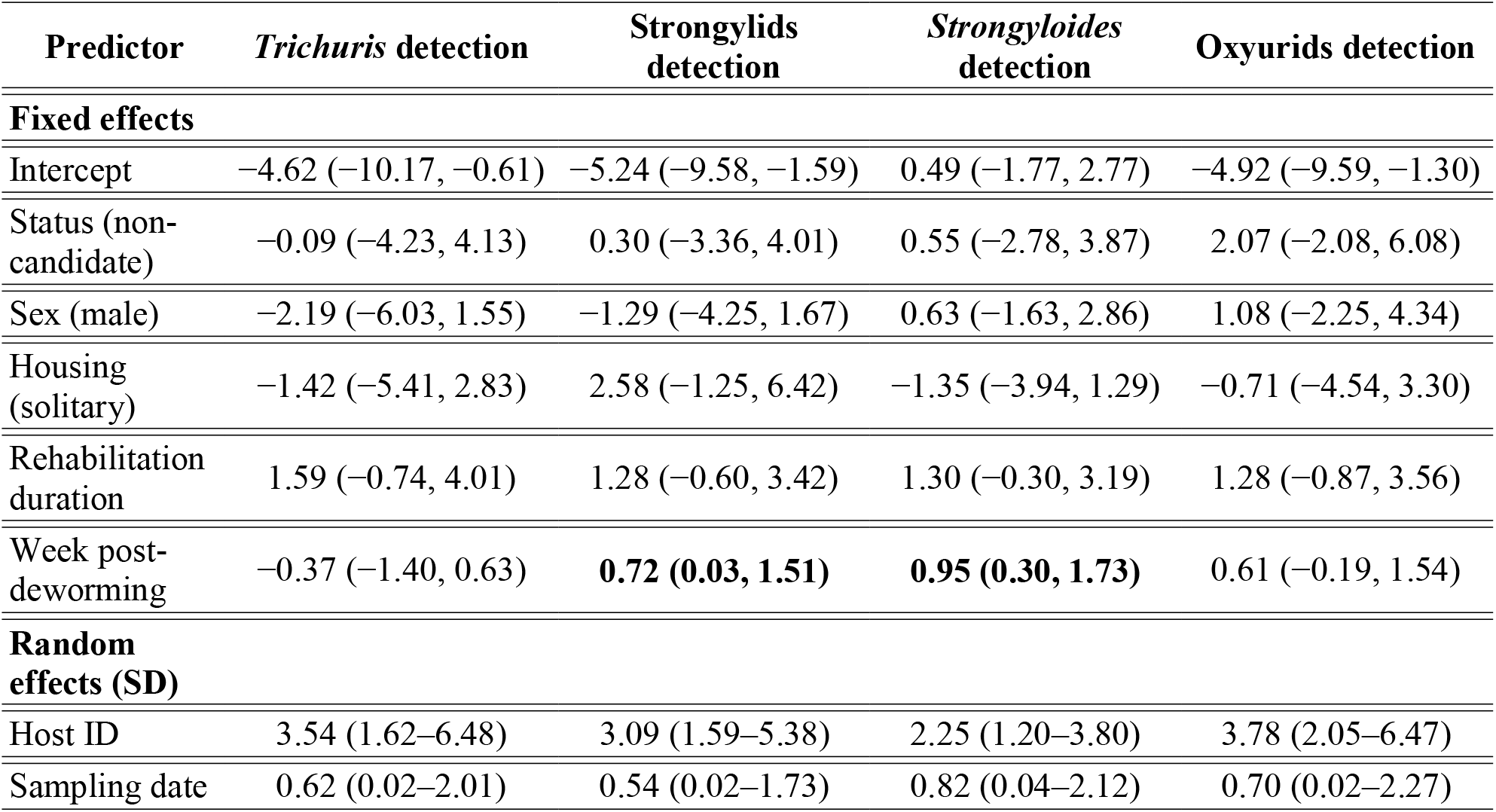
Bayesian mixed-effects logistic regression results for helminth infection probability in rehabilitating *N. javanicus*. Posterior means (β) and 95% credible intervals (CrI).Posterior means (β) and 95% credible intervals (CrI).

**Figure 1.**
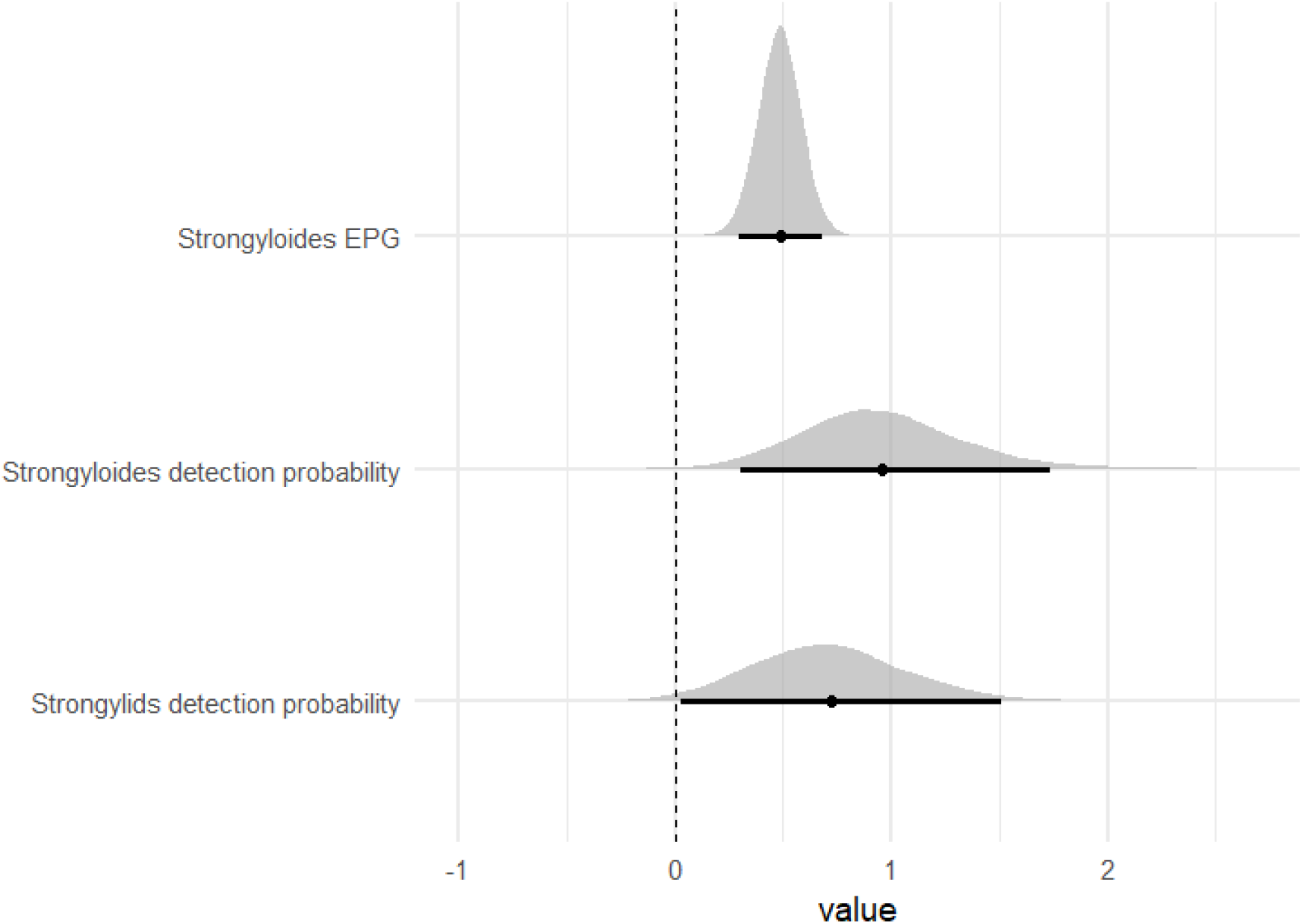
Posterior distributions of the effect of weeks post-deworming for parasite models showing credible temporal associations.

### Parasite intensity

Bayesian negative binomial mixed-effects models revealed limited support for host traits or management-related variables on helminth egg counts. Among the four parasite groups examined, only *Strongyloides* spp. exhibited a credible positive association between EPG and weeks post-deworming, indicating increasing egg counts over time following treatment (Figure 1; Figure S5). For *Trichuris* spp., strongylids, and oxyurids, no strong evidence was found for effects of release status, sex, housing condition, or weeks post-deworming on egg counts, with all 95% credible intervals overlapping zero. Across all models, substantial individual-level variation in egg counts was evident, as reflected by the random-effect variance associated with host identity (Table 3).

**Table 3.** Bayesian negative binomial mixed-effects models results for helminth egg counts (EPG) in rehabilitating *N. javanicus*. Posterior means (β) and 95% credible intervals (CrI).

| Predictor | *Trichuris* EPG | Strongylids EPG | *Strongyloides* EPG | Oxyurids EPG |
| --- | --- | --- | --- | --- |
| **Fixed effects** |  |  |  |  |
| Intercept | −4.85 (−9.75, −1.07) | −2.28 (−5.60, 0.62) | 3.60 (1.57, 5.51) | −3.13 (−6.97, 0.18) |
| Status (non-candidate) | 0.26 (−1.67, 2.11) | 0.60 (−1.28, 2.42) | 0.87 (−0.83, 2.47) | 0.72 (−1.11, 2.56) |
| Sex (male) | −0.47 (−2.37, 1.43) | −0.23 (−2.04, 1.57) | 0.53 (−1.06, 2.08) | 0.27 (−1.61, 2.13) |
| Housing (solitary) | −0.08 (−2.02, 1.82) | 0.69 (−1.14, 2.59) | −0.51 (−2.13, 1.25) | 0.12 (−1.74, 2.00) |
| Week post-deworming | 0.16 (−0.27, 0.54) | 0.27 (−0.00, 0.54) | **0.49 (0.29, 0.68)** | 0.16 (−0.11, 0.42) |
| **Random effects (SD)** |  |  |  |  |
| Host ID | 5.44 (3.40–8.45) | 5.06 (3.19–7.72) | 2.86 (1.83–4.30) | 5.64 (3.63–8.58) |
| Sampling date | 0.33 (0.01–0.89) | 0.19 (0.01–0.53) | 0.11 (0.00–0.32) | 0.21 (0.01–0.60) |

## DISCUSSION

This study demonstrated that gastrointestinal helminth infection patterns in rehabilitating *N. javanicus* are largely not explained by host classification or rehabilitation status based on general health and medical history. Host traits and management categories, including release candidacy, sex, and housing condition, explained little of the observed variation. Where temporal associations were detected, these were restricted to a subset of parasite taxa: detection probability increased with time since anthelmintic treatment for strongylids and *Strongyloides* spp., and infection intensity also increased over time for *Strongyloides* spp. alone. Parasite richness and community diversity remained relatively stable among individuals. The residual variation was characterised in part by post-treatment temporal dynamics in these select taxa, alongside substantial individual-level heterogeneity.

### Temporal dynamics dominate helminth infection patterns during rehabilitation

The association between time since anthelmintic treatment and detection probability in strongylids and *Strongyloides* spp. is consistent with the fundamentally dynamic nature of helminth infections in rehabilitation settings. Rather than reflecting fixed attributes of host condition, these patterns suggest that parasite infections in captive primates are shaped by cycles of treatment, reinfection, and environmental exposure(Grove et al., 1988; Altizer et al., 2006; Pedersen and Fenton, 2007), with veterinary interventions modulating infection intensity transiently rather than eliminating infection risk.

*Strongyloides* and strongylids, which have direct life cycles and persist as infective stages in captive environments, are particularly prone to rapid reinfection following treatment (Grove et al., 1988; Schapiro and Bernacky, 2012; Li et al., 2015). The post-treatment increases in infection probability and intensity observed for these taxa most likely reflect reinfection dynamics, though incomplete treatment efficacy cannot be excluded. Albendazole performs poorly against *Trichuris* spp., which may account for the absence of a temporal signal in that group (Patel et al., 2020; Gebreyesus et al., 2024). For oxyurids and strongylids, although albendazole efficacy is relatively high (∼94% and ∼92%) (Wendt et al., 2019; Horton, 2000), the absence of a detectable temporal signal most likely reflects low overall prevalence in our samples, which limits statistical power to detect post-treatment dynamics even if they occur, rather than a genuine absence of treatment or reinfection effects. Similar treatment-linked temporal patterns have been reported across managed primate populations, where standardized veterinary interventions reduce infection intensity transiently but do not prevent reinfection (Stringer and Linklater, 2014; Herrera et al., 2019). These findings reinforce the view that anthelmintic treatment is best understood as a tool for managing infection burden rather than achieving parasite clearance, and that ongoing monitoring is necessary given the inevitability of reinfection in non-sterile captive environments.

### Individual heterogeneity outweighs coarse host classifications

Despite clear differences in rehabilitation history and release candidacy status, parasite richness, diversity, detection probability, and abundance showed substantial overlap among individuals. Random-effect variance associated with host identity accounted for a large proportion of unexplained variation, indicating pronounced individual-level heterogeneity in parasite burdens. Such heterogeneity is a common feature of host–parasite systems and reflects variation in exposure, susceptibility, immune function, and stochastic infection processes that are not readily captured by coarse categorical predictors (Bush et al., 1997; Pedersen and Fenton, 2007).

The absence of strong effects of sex, housing condition, or rehabilitation duration aligns with previous studies of slow lorises and other captive primates, where expected host-based predictors often fail to show consistent associations with helminth infection under managed conditions (Li et al., 2015; Rode-Margono et al., 2015; Herrera et al., 2019). A broader comparison across rehabilitated and captive primate studies further indicates that parasite prevalence, dominant taxa, and inferred risk factors vary widely among facilities and study systems, even under ostensibly similar management regimes (Table S6), underscoring the dominance of context-dependent and individual-level processes over coarse host classifications. Standardized husbandry, uniform veterinary care, and controlled diets may homogenize exposure pathways and partially override differences arising from individual health histories. These findings are consistent with broader evidence that condition–infection relationships in wildlife are highly variable and context dependent, with null effects being common rather than exceptional (Sanchez et al., 2018). This interpretation is further supported by a general synthesis emphasizing that individual-level heterogeneity, rather than coarse host classifications, often dominates wildlife disease dynamics and responses to management interventions (McDonald et al., 2018).

### Limits of parasite metrics as indicators of release readiness

A key implication of our results is that gastrointestinal parasite metrics do not clearly correspond to existing classifications of release readiness in rehabilitating *N. javanicus*. Although rehabilitation duration was not retained as a statistically definitive predictor under a strict credible-interval interpretation, posterior effect sizes suggested that longer time in rehabilitation may be associated with increased odds of infection detection for some parasite taxa. Release candidates defined on the basis of physical condition and behavioral criteria did not exhibit consistently lower parasite prevalence or abundance than non-candidates, despite having shorter rehabilitation histories on average. This finding cautions against interpreting parasite burdens as direct proxies for host quality or readiness for translocation. Instead, parasites should be considered part of the normal host-associated biota, with substantial variation observed among individuals. This individual-level heterogeneity suggests that parasite dynamics may reflect host-specific processes and reinforces the value of longitudinal monitoring approaches that track changes within individuals over time. Future work is therefore needed to evaluate how infection dynamics during rehabilitation relate to host health and post-release outcomes.

Rather, parasite infections in rehabilitation contexts appear to reflect recent treatment history and environmental exposure more strongly than broad differences in host classification,consistent with our finding that time since anthelmintic treatment was the main detectable predictor of infection in some parasite taxa. Similar concerns have been raised in other rehabilitation and translocation programs, where reliance on single health metrics canoversimplify biologically complex and temporally dynamic processes (Altizer et al., 2006; Herrera et al., 2019). From this perspective parasite data are best interpreted as indicators of ongoing host-parasite interactions within rehabilitation environments rather than as fixed thresholds for release decisions(Cope et al., 2022).

### Implications for parasite management in rehabilitation programs

From a management perspective, the temporal dynamics and individual-level variation in parasite burden observed here underscore the importance of continuous monitoring rather than one-time assessment. While standardized veterinary care and routine anthelmintic treatment are effective in limiting chronic parasite burdens, they do not eliminate reinfection risk, particularly for directly transmitted helminths such as *Strongyloides* (Schapiro and Bernacky, 2012; Li et al., 2015; Hildebrand and Zalesny, 2025). Moreover, treatment efficacy likely varies across parasite taxa. Albendazole, for instance, is known to be less effective against *Trichuris* spp., suggesting that adaptive treatment protocols accounting for taxon-specific resistance patterns may be more effective than standardized regimens applied uniformly across individuals. This supports treating parasite management as an iterative and adaptive process throughout captivity rather than a discrete pre-release intervention.

Integrating consistent parasite monitoring with other health and behavioral assessments may therefore provide a more robust framework for managing infection risks during rehabilitation and minimizing potential impacts on post-release survival or pathogen spillover into wild populations. More broadly, recent syntheses emphasize that combining parasite dynamics with complementary indicators such as behavior and host-associated microbiomes can improve conservation-relevant health assessments by capturing biological processes operating across different temporal scales (Cope et al., 2022). Such integrative approaches are increasingly feasible in rehabilitated primates, where behavioral and microbiome data can be collected alongside parasitological monitoring, as suggested by recent work on rehabilitating slow lorises (Langgeng et al., 2026).

## Acknowledgements

We would like to thank Kyoto University Wildlife Research Center (WRC), CICASP, Research Units for Exploring Future Horizons (Coevolution and Coexistence), and the Joint Research Program of WRC for facilitating our research, YIARI Bogor board of directors and staff (especially Mastur, Ajo, Aconk, Ganyong, Pak Aki, Kojek) for their permission and assistance in the field, FMIPA-Biologi IPB University dean and staff. We also thanked BRIN, KLHK, Dirjen KSDAE, and BBKSDAE Jawa Barat for the administrative assistance and permission to conduct research in Indonesia.

## Author contributions

AL: Conceptualization, formal analysis, methodology, investigation, writing – original draft, funding acquisition, project administration. WP: writing – review & editing, project administration, investigation. NPP: writing – review & editing, project administration, investigation. PR: writing – review & editing, project administration. RM: writing – review & editing, project administration. MS: conceptualization, writing – review & editing, supervision, funding acquisition. AJJM: conceptualization, writing – review & editing, formal analysis, supervision, funding acquisition. IM: conceptualization, formal analysis, writing – original draft, supervision, funding acquisition

## Data availability

All data needed to evaluate the conclusions in the paper are present in the paper. Additional data related to this paper may be requested from the authors.

## Declarations

### Ethical approval

This research was conducted in accordance with the Guidelines for the Care and Use of Non-human Primates and the Guidelines for Field Research on Non-human Primates established by the Center for the Evolutionary Origins of Human Behavior, Kyoto University. Ethical approval and permissions were obtained from the Field Research Committee of the Wildlife Research Center, Kyoto University, as well as the Indonesian National Research and Innovation Agency (BRIN, Ref. no.: 009/KE.02/SK/01/2024). Permission to collect biological samples from nationally protected species were obtained from the Director General of Natural Resources and Ecosystem Conservation (Dirjen KSDAE, Ref. no.: SK.127/KSDAE/SETKSDAE/KSA2/6/2024).

### Generative AI and AI-assisted technologies in the writing process

During the preparation of this manuscript, the authors used ChatGPT-5.2 to refine the clarity and logical flow of the text. The authors carefully reviewed, corrected, and approved all content generated, and take full responsibility for the final published version.

