## Supplementary Materials for "Helminth infection dynamics in rehabilitating Javan slow lorises are driven by time since deworming rather than host traits"

Table S1. Study subject and fecal sampling summary for rehabilitating *Nycticebus javanicus* included in this study. Shown are individual identity, sex, release candidacy status (candidate or non-candidate), housing condition (group or solitary), year of admission to the YIARI rehabilitation center, and the number of fecal samples collected prior to deworming or for non-release candidates (“Pre/non”) and during the soft-release period for release candidates (“Soft”). All individuals were adults at the time of sampling.

| **ID** | **Sex** | **Release candidacy status** | **Housing condition** | **Year admitted to YIARI** | **Number of fecal samples collected** | |
| --- | --- | --- | --- | --- | --- | --- |
|  |  |  |  |  | Pre/non | Soft |
| **JS01** | Female | Candidate | Solitary | 2022 | 13 | 1 |
| **JS02** | Female | Candidate | Group | 2024 | 6 | 0 |
| **JS03** | Female | Candidate | Solitary | 2023 | 12 | 1 |
| **JS04** | Male | Candidate | Solitary | 2024 | 11 | 1 |
| **JS05** | Female | Candidate | Group | 2022 | 8 | 0 |
| **JS06** | Female | Candidate | Group | 2024 | 10 | 1 |
| **JS07** | Female | Candidate | Solitary | 2023 | 10 | 1 |
| **JS08** | Female | Candidate | Group | 2024 | 6 | 1 |
| **JS09** | Male | Candidate | Solitary | 2024 | 12 | 1 |
| **JS10** | Male | Candidate | Group | 2024 | 5 | 0 |
| **JS11** | Female | Non-candidate | Solitary | 2016 | 6 | 0 |
| **JS12** | Female | Non-candidate | Solitary | 2012 | 5 | 0 |
| **JS13** | Male | Non-candidate | Solitary | 2017 | 6 | 0 |
| **JS14** | Female | Non-candidate | Solitary | 2022 | 5 | 0 |
| **JS15** | Male | Non-candidate | Solitary | 2015 | 5 | 0 |
| **JS16** | Female | Non-candidate | Solitary | 2015 | 4 | 0 |
| **JS17** | Male | Non-candidate | Solitary | 2022 | 4 | 0 |
| **JS18** | Male | Non-candidate | Solitary | 2018 | 5 | 0 |
| **JS19** | Female | Non-candidate | Solitary | 2018 | 7 | 0 |

Table S2. Pareto-smoothed importance sampling leave-one-out cross-validation (PSIS-LOO) diagnostics for Bayesian models of parasite diversity metrics in rehabilitating *N. javanicus*. The expected log predictive density (elpd_loo) summarizes predictive performance, p_loo represents the effective number of parameters, and LOOIC is the leave-one-out information criterion. Pareto-k diagnostics indicate that nearly all observations fell within the reliable range (k ≤ 0.7), suggesting stable cross-validation estimates.

| Model | elpd_loo (SE) | p_loo (SE) | LOOIC (SE) | k ≤ 0.7 | 0.7 < k ≤ 1 | k > 1 |
| --- | --- | --- | --- | --- | --- | --- |
| Shannon diversity | 24.36 (14.81) | 23.03 (4.89) | −48.71 (29.62) | 146 (99.3%) | 1 (0.7%) | 0 (0.0%) |
| Parasite richness | −164.97 (6.53) | 6.15 (0.43) | 329.95 (13.07) | 147 (100%) | 0 (0.0%) | 0 (0.0%) |

Table S3. Pareto-smoothed importance sampling leave-one-out cross-validation (PSIS-LOO) diagnostics for Bayesian Bernoulli mixed-effects models of parasite detection probability. The expected log predictive density (elpd_loo) summarizes model predictive performance, p_loo represents the effective number of parameters, and LOOIC is the leave-one-out information criterion. Pareto-k diagnostics indicate that the majority of observations fell within the reliable range (k ≤ 0.7), suggesting stable cross-validation estimates.

| Model | elpd_loo (SE) | p_loo (SE) | LOOIC (SE) | k ≤ 0.7 | 0.7 < k ≤ 1 | k > 1 |
| --- | --- | --- | --- | --- | --- | --- |
| Trichuris prevalence | −16.98 (3.45) | 5.19 (1.29) | 33.97 (6.91) | 146 (99.3%) | 1 (0.7%) | 0 (0.0%) |
| Strongylids prevalence | −41.27 (5.61) | 12.01 (2.25) | 82.55 (11.21) | 144 (98.0%) | 3 (2.0%) | 0 (0.0%) |
| Strongyloides prevalence | −78.95 (6.15) | 25.58 (2.61) | 157.89 (12.30) | 146 (99.3%) | 1 (0.7%) | 0 (0.0%) |
| Oxyurids prevalence | −28.84 (5.33) | 11.29 (2.64) | 57.69 (10.66) | 144 (98.0%) | 3 (2.0%) | 0 (0.0%) |

Table S4. Odds ratios (OR) and 95% credible intervals (CrI) for parasite detection probability models

| Predictor | *Trichuris* detection OR (95% CrI) | Strongylids detection OR (95% CrI) | *Strongyloides* detection OR (95% CrI) | Oxyurids detection OR (95% CrI) |
| --- | --- | --- | --- | --- |
| Status (non-candidate) | 0.91 (0.01–61.56) | 1.35 (0.03–55.15) | 1.73 (0.06–47.94) | 7.92 (0.13–436.84) |
| Sex (male) | 0.11 (0.00–4.71) | 0.28 (0.01–5.31) | 1.88 (0.20–17.46) | 2.94 (0.11–76.66) |
| Housing (solitary) | 0.24 (0.00–16.95) | 13.20 (0.29–613.11) | 0.26 (0.02–3.63) | 0.49 (0.01–27.11) |
| Rehabilitation duration | 4.90 (0.48–55.15) | 3.60 (0.55–30.57) | 3.67 (0.74–24.29) | 3.60 (0.42–35.16) |
| Weeks post-deworming | 0.69 (0.25–1.88) | 2.05 (1.03–4.53) | 2.59 (1.35–5.64) | 1.84 (0.83–4.66) |

Table S5. Pareto-smoothed importance sampling leave-one-out cross-validation (PSIS-LOO) diagnostics for Bayesian zero-inflated negative binomial models of parasite egg counts in rehabilitating *N. javanicus*. The expected log predictive density (elpd_loo) summarizes predictive accuracy, while p_loo represents the effective number of parameters. Pareto-k diagnostics indicate that the majority of observations fell within the reliable range (k ≤ 0.7), suggesting stable cross-validation estimates.

| Model | elpd_loo (SE) | p_loo (SE) | LOOIC (SE) | k ≤ 0.7 | 0.7 < k ≤ 1 | k > 1 |
| --- | --- | --- | --- | --- | --- | --- |
| Trichuris EPG | −104.46 (24.39) | 6.59 (2.09) | 208.92 (48.77) | 145 (98.6%) | 1 (0.7%) | 1 (0.7%) |
| Strongylids EPG | −206.11 (29.59) | 10.60 (2.08) | 412.22 (59.17) | 143 (97.3%) | 3 (2.0%) | 1 (0.7%) |
| Strongyloides EPG | −541.16 (39.08) | 20.11 (3.01) | 1082.33 (78.16) | 144 (98.0%) | 3 (2.0%) | 0 (0.0%) |
| Oxyurids EPG | −176.37 (28.56) | 8.60 (1.74) | 352.74 (57.12) | 142 (96.6%) | 5 (3.4%) | 0 (0.0%) |

Table S6. Published records of gastrointestinal helminths infecting slow lorises (*Nycticebus* spp.) across wild, captive, confiscated, and rehabilitation contexts. The table summarizes reported helminth taxa, host species, study location, and management context based on available literature and husbandry manuals. Records include both identified parasite species and higher taxonomic or unidentified groups, reflecting variation in diagnostic resolution among studies.

| **Parasite Taxon** | **Parasite Species/Type** | **Host Species** | **Location/Study Type** | **Reference** |
| --- | --- | --- | --- | --- |
| **NEMATODA** |  |  |  |  |
| **Oxyuridae** | *Lemuricola* (*Protenterobius*) *nycticebi* | *N. javanicus* | Wild, West Java, Indonesia | Rode-Margono et al. (2015) |
| **Oxyuridae** | *Lemuricola* (*Protenterobius*) *nycticebi* | *N. bengalensis* | Captive/confiscated, Bangladesh | Mondal et al. (2026) |
| **Oxyuridae** | *Oxyuris* spp. | *N. coucang* | Captive, YIARI, Indonesia | Ulfa (2014) |
| **Oxyuridae** | *Enterobius* spp. | *N. pygmaeus* | Captive, rescue center | Streicher (2004) cited in Rode-Margono et al. (2015) |
| **Oxyuridae** | *Enterobius* spp. | *N. pygmaeus* | Captive, Duke Primate Center, USA | San Diego Zoo Loris Husbandry Manual |
| **Oxyuridae** | Oxyurids (unidentified) | *N. coucang* | Captive, Duke Primate Center, USA | San Diego Zoo Loris Husbandry Manual |
| **Oxyuridae** | Oxyurids (unidentified) | *N. javanicus* | Captive, YIARI, Indonesia | Wibowo (2014) |
| **Ascarididae** | *Ascaris* spp. | *N. coucang* | Captive, YIARI, Indonesia | Ulfa (2014) |
| **Ascarididae** | *Ascaris* spp. | *N. javanicus* | Captive, YIARI, Indonesia | Wibowo (2014) |
| **Ascarididae** | *Ascaris* spp. | *N. coucang* | Captive, literature review | Setyorini & Werdateti (2005) |
| **Ancylostomatidae** | *Necator* spp. (hookworm) | *N. javanicus* | Wild, West Java, Indonesia | Rode-Margono et al. (2015) |
| **Strongyloididae** | *Strongyloides* spp. | *N. coucang* | Captive, YIARI, Indonesia | Ulfa (2014) |
| **Strongyloididae** | *Strongyloides* spp. | *N. javanicus* | Captive, YIARI, Indonesia | Wibowo (2014) |
| **Strongyloididae** | *Strongyloides* sp. | *N. menagensis* | Wild, Malaysian Borneo | Frias et al. (2018) |
| **Strongyloididae** | *Strongyloides* sp. | *N. coucang* | Captive, Duke Primate Center, USA | San Diego Zoo Loris Husbandry Manual |
| **Strongylidae** | Strongylids (unidentified) | *N. coucang* | Captive, YIARI, Indonesia | Ulfa (2014) |
| **Strongylidae** | Strongylids (unidentified) | *N. javanicus* | Captive, YIARI, Indonesia | Wibowo (2014) |
| **Trichostrongylidae** | *Trichostrongylus* spp. | *N. javanicus* | Wild, West Java, Indonesia | Rode-Margono et al. (2015) |
| **Trichuridae** | *Trichuris* spp. | *N. coucang* | Captive, YIARI, Indonesia | Ulfa (2014) |
| **Trichuridae** | *Trichuris* spp. | *N. pygmaeus* | Captive, San Diego Zoo, USA | San Diego Zoo Loris Husbandry Manual |
| **Molineidae** | *Pterygodermatites nycticebi* | *N. coucang* | Captive, San Diego Zoo, USA | San Diego Zoo Loris Husbandry Manual |
| **Physalopteridae** | *Physaloptera* sp. | *N. coucang* | Captive, Duke Primate Center, USA | San Diego Zoo Loris Husbandry Manual |
| **Spiruridae (unidentified)** | Nematodes (unidentified) | *N. coucang* | Captive, San Diego Zoo, USA | San Diego Zoo Loris Husbandry Manual |
| **CESTODA** |  |  |  |  |
| **Hymenolepididae** | *Hymenolepis* spp. | *N. coucang* | Captive, YIARI, Indonesia | Ulfa (2014) |
| **Hymenolepididae** | *Hymenolepis* sp. | *N. javanicus* | Captive, YIARI, Indonesia | Wibowo (2014) |
| **Hymenolepididae** | *Hymenolepis*-like ova | *N. pygmaeus* | Captive, San Diego Zoo, USA | San Diego Zoo Loris Husbandry Manual |
| **Anoplocephalidae** | Tapeworms (unidentified) | *N. coucang* | Captive, San Diego Zoo, USA | San Diego Zoo Loris Husbandry Manual |

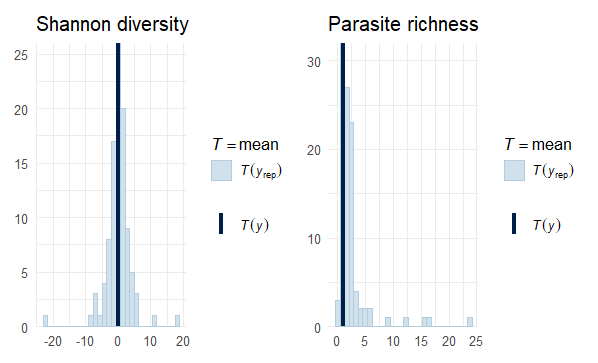

Figure S1. Prior predictive checks for parasite Shannon diversity and richness models. Histograms show the distribution of simulated mean values under the prior predictive distribution; vertical lines indicate the observed mean. In both cases, the observed values fell within plausible prior predictive ranges.

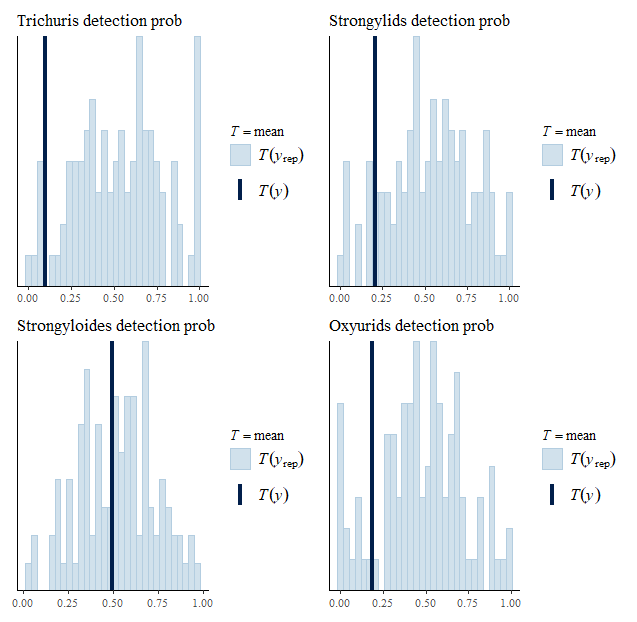

Figure S2. Prior predictive checks for parasite detection probability. Histograms show the distribution of simulated mean values under the prior predictive distribution; vertical lines indicate the observed mean. In all cases, the observed values fell within plausible prior predictive ranges.

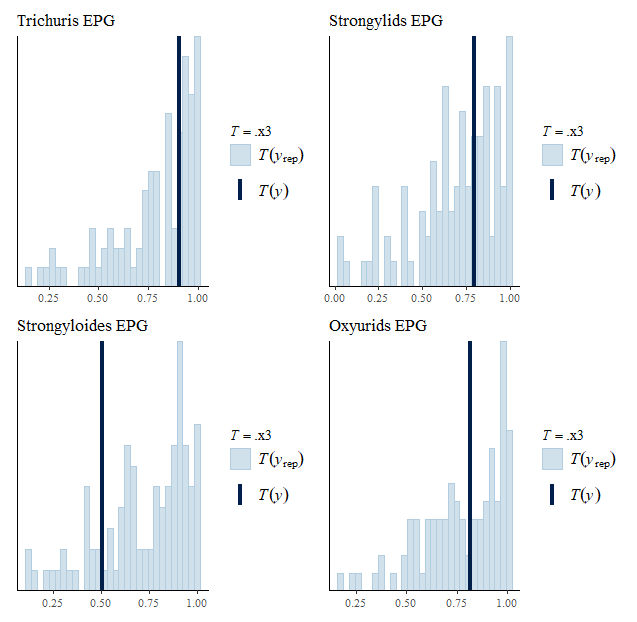

Figure S3. Prior predictive checks for parasite EPG models. Histograms show the distribution of simulated mean values under the prior predictive distribution; vertical lines indicate the observed mean. In all cases, the observed values fell within plausible prior predictive ranges.

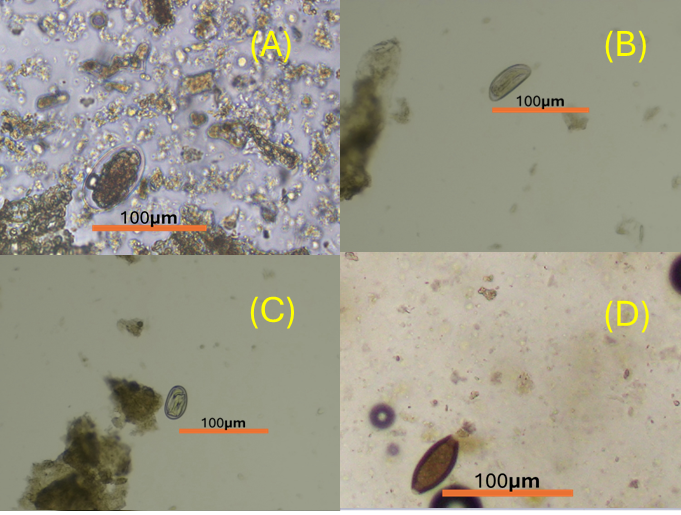

Figure S4. GI helminth eggs infecting rehabilitating *N. javanicus*: (A) Strongylids (B) Oxyurids, (C) *Strongyloides*, (D) *Trichuris*

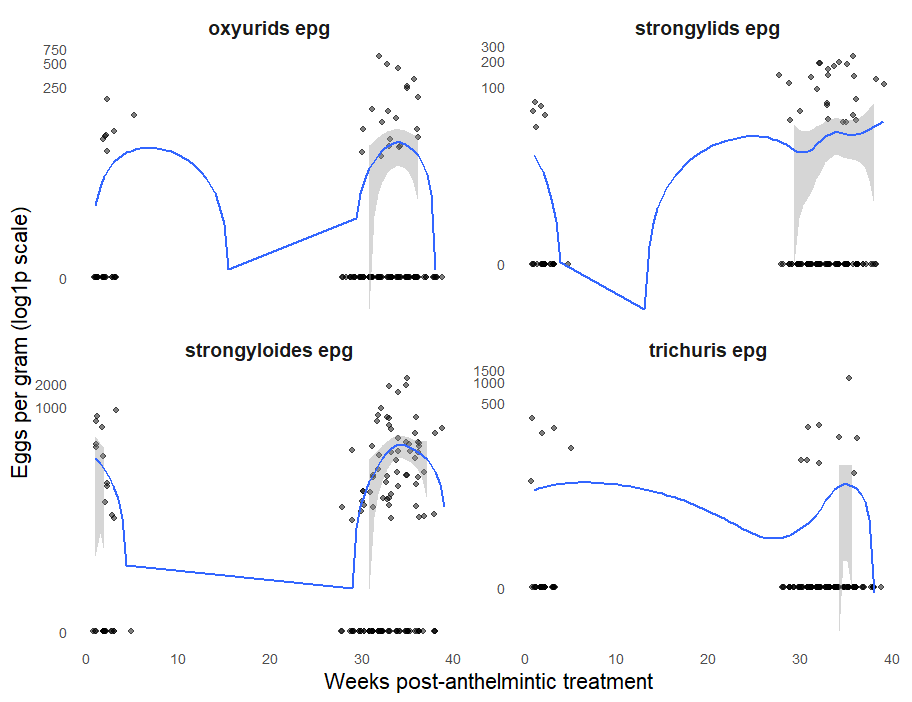

Figure S5. Density plot of parasite EPG related to weeks post-anthelmintic treatment
